# Telomere-to-telomere, gap-free genome of mung beans (*Vigna radiata*) provides insights into domestication under structural variation

**DOI:** 10.1101/2024.08.21.609077

**Authors:** Kai-Hua Jia, Guan Li, Longxin Wang, Min Liu, Zhi-Wei Wang, Ru-Zhi Li, Lei-Lei Li, Kun Xie, Yong-Yi Yang, Ru-Mei Tian, Xue Chen, Yu-Jun Si, Xiao-Yan Zhang, Feng-Jing Song, Na-Na Li

## Abstract

Mung bean (*Vigna radiata*), an essential annual legume, holds substantial value in global agriculture due to its short growth cycle, low input requirements, and nutritional benefits. Despite extensive domestication, the genetic mechanisms underlying its morphological and physiological evolution remain incompletely understood. In this study, we present a gap-free, telomere-to-telomere genome assembly of the mung bean cultivar ‘Weilv-9’, achieved through the integration of PacBio HiFi, Oxford Nanopore, and Hi-C sequencing technologies. The 500 Mb assembly, encompassing 11 chromosomes and containing 28,740 protein-coding genes, reveals that 49.17% of the genome comprises repetitive sequences. Within the genome, we found the recent amplification of transposable elements significantly impacts the expression of nearby genes. Furthermore, integrating structural variation and SNP data from resequencing, we identified that the fatty acid synthesis, suberin biosynthetic, and phenylpropanoid metabolic processes have undergone strong selection during domestication. These findings provide valuable insights into the genetic mechanisms driving domestication and offer a foundation for future genetic enhancement and breeding programs in mung beans and related species.

## Introduction

Mung bean (*Vigna radiata*), also known as Asia’s Ancient Bean, is a significant annual, warm-season legume crop, accounting for approximately 8.5% of the global legume cultivation area, with over 7.3 million hectares cultivated worldwide (Nair & Schreinemachers, 2020). India, Myanmar, and China are the primary producers. This crop is extensively grown in Asian countries (over 90%) and in the warmer regions of western countries due to its favorable attributes, such as a short growth cycle (70–90 days), low input requirements, drought tolerance, and nitrogen fixation capabilities (Ha & Lee, 2019). Mung bean seeds are nutritionally rich, containing a balanced profile of macro– and micro-nutrients, including proteins, dietary fibers, vitamins, and minerals, as well as high levels of bioactive components (Hou *et al*., 2019). Traditionally, mung beans have been consumed with cereals in China and India to complement the lysine deficiency in cereal grains. Nowadays, they are commonly used in fresh, boiled, processed, and canned food products. Their high digestible protein content makes them particularly suitable for vegetarian and vegan diets. Additionally, mung beans are an excellent protein source for children due to their low allergenic and flatulence-causing components (Ha & Lee, 2019). The increasing awareness of health and the prevalence of lifestyle-related diseases globally have driven the growing demand for mung beans.

The domestication history of mung beans can be traced back to its wild progenitor, *V. radiata* var. *sublobata*, approximately 50,000 years ago, in the tropical and subtropical regions of the South Asian subcontinent (Lin *et al*., 2023). Despite the extensive domestication process, leading to significant morphological and physiological differences between cultivated and wild types, there are virtually no reproductive barriers between them, which endows mung beans with a high degree of genetic diversity and potential for improvement (Ignacimuthu & Babu, 1987; Pandiyan *et al*., 2010; Thuan, 2011). Whole-genome resequencing, particularly targeting SNP (single nucleotide polymorphism) variation sites, presents the most direct approach for studying domestication (Xu & Bai, 2015). This method allows for a comprehensive analysis of genetic diversity and the identification of domestication-related loci. By comparing domesticated species with their wild relatives, researchers can pinpoint specific genetic changes that have occurred during domestication processes (Nan *et al*., 2022; Wu *et al*., 2023). Furthermore, SNP variation analysis provides insights into evolutionary dynamics, selective sweeps, and genetic bottlenecks associated with domestication events.

While SNPs have significantly advanced our understanding of domestication, structural variations (SVs), including insertions (INSs), deletions (DELs), inversions (INVs), duplications (DUP), and translocations (TRAs), offer an additional layer of genetic complexity that is equally crucial for understanding the full spectrum of genetic changes associated with domestication (Hitte, 2019; Cumer *et al*., 2021). SVs can have a more profound impact on genome structure and function than SNPs, as they often involve larger segments of DNA and can affect multiple genes or regulatory regions simultaneously. For instance, in humans, the likelihood of SVs being associated with phenotypic traits is threefold higher than that of SNPs (Chaisson *et al*., 2019). They also play a vital role in domestication processes. In the context of domestication, SVs are identified as the causative genetic variants for at least one-third of known domestication alleles (Gaut *et al*., 2018). Numerous instances demonstrate that SV is a key driver of domestication. For example, in tomatoes, multiple SVs have been found to modify gene dosage and expression levels, thereby influencing fruit flavor, size, and yield (Alonge *et al*., 2020). In grapes, several SVs have been involved in coastal adaptation (Zhang *et al*., 2024). In soybeans, SVs have impacted the domestication of genes involved in the fatty acid biosynthesis pathway (Jia *et al*., 2024). However, detailed studies on the patterns of SV variation during mung bean domestication are still limited.

Recent technological advancements in genome sequencing have facilitated the development of high-resolution reference genomes for mung bean (Kang *et al*., 2014; Ha *et al*., 2021; Liu *et al*., 2022). However, due to the prevalence of highly tandem repetitive sequences in genomes, such as centromeric, ribosomal DNA (rDNA), and transposable element (TE) sequences, achieving complete telomere-to-telomere (T2T) genome assembly remains challenging. In this research, we produced comprehensive T2T and gap-free genome assemblies for the cultivated mung bean “Weilv-9“ employing an integrated approach of Pacific Biosciences (PacBio), Oxford Nanopore Technologies (ONT), and high throughput chromosome conformation capture (Hi-C) sequencing technologies. The meticulous assemblies span 500 Mb in length, encapsulating 28,740 protein-encoding genes. By comparing with genomic data from cowpea and common bean, we discovered that recently amplified TEs in mung bean significantly altered the expression of nearby genes. Utilizing these T2T genome assemblies along with 34 wild and 77 cultivated high-depth resequenced mung bean datasets, comprehensive analyses integrating both SNP and SV data identified potential domestication genes linked to plant architecture transformation. Collectively, this refined mung bean genome and SVs offer unparalleled insight into the genomic shifts underpinning mung bean evolution and domestication, establishing a cornerstone for future mung bean investigations and cultivar development.

## Results

### Gap-free T2T genome assembly of mung bean

The size of the mung bean genome has previously been estimated or assembled to range from 475 Mb to 484 Mb (Kang *et al*., 2014; Ha *et al*., 2021; Liu *et al*., 2022). We have chosen to perform genome sequencing on “Weilv-9”, a mung bean variety characterized by its high yield and stress resistance, along with a compact plant form. In order to assemble the accurate and complete these genomes possible, we selected a hybrid sequencing approach. This included PacBio High-Fidelity (HiFi) long reads (46 Gb), ONT ultra-long reads (53 Gb) and Hi-C paired-end reads (64 Gb) (**Supplementary Table 1**).

In our initial approach, we employed the *de novo* assembler HiFiasm (Cheng *et al*., 2021) to process HiFi, ONT, and Hi-C reads for contig generation. Subsequently, the Hi-C reads were aligned to the primary assembly utilizing Juicer (Durand *et al*., 2016). This step was followed by a preliminary chromosome assembly aided by Hi-C, using 3d-DNA (Dudchenko *et al*., 2017). Manual verification and adjustments were conducted on the alignments within Juicebox (Robinson *et al*., 2018) to ensure accuracy and correct orientation of sequences. Further refinement involved the use of LR_Gapcloser (Xu *et al*., 2019) to align ONT reads to the genome, specifically targeting bridging reads that spanned existing gaps. This process facilitated the integration of ONT read sequences into genomic gaps. Additionally, we undertook the reassembly of HiFi reads aligned at scaffold ends, aiming to extend their lengths. In parallel, the complete chloroplast and mitochondrial genomes were assembled using GetOrganelle (Jin *et al*., 2020). The culmination of these methodologies resulted in the production of comprehensive and gap-free mung bean genome assemblies (**Fig. 1**). These assemblies encompassed 11 chromosomes along with mitochondrial and chloroplast genomes, collectively spanning a total length of 500 Mb (**Table 1**). The contig/scaffold N50 of the mung bean genome were 45 Mb. Through searching for the canonical ‘TTTAGGG/CCCTAAA’ telomeric repeat, we identified 19 the telomeres, with CCCTAAA at the 5’ end and TTTAGGG at the 3’ end (**Supplementary Fig. 1A**). The distribution of the 18-5.8-28S rDNA is primarily localized at the 5’ end of chromosome 1 and the 3’ end of chromosome 11 (**Supplementary Fig. 1B**). Conversely, the 5S rDNA shows a predominant presence on chromosomes 3, 4, and 8 (**Supplementary Fig. 1C**).

**Fig. 1.**
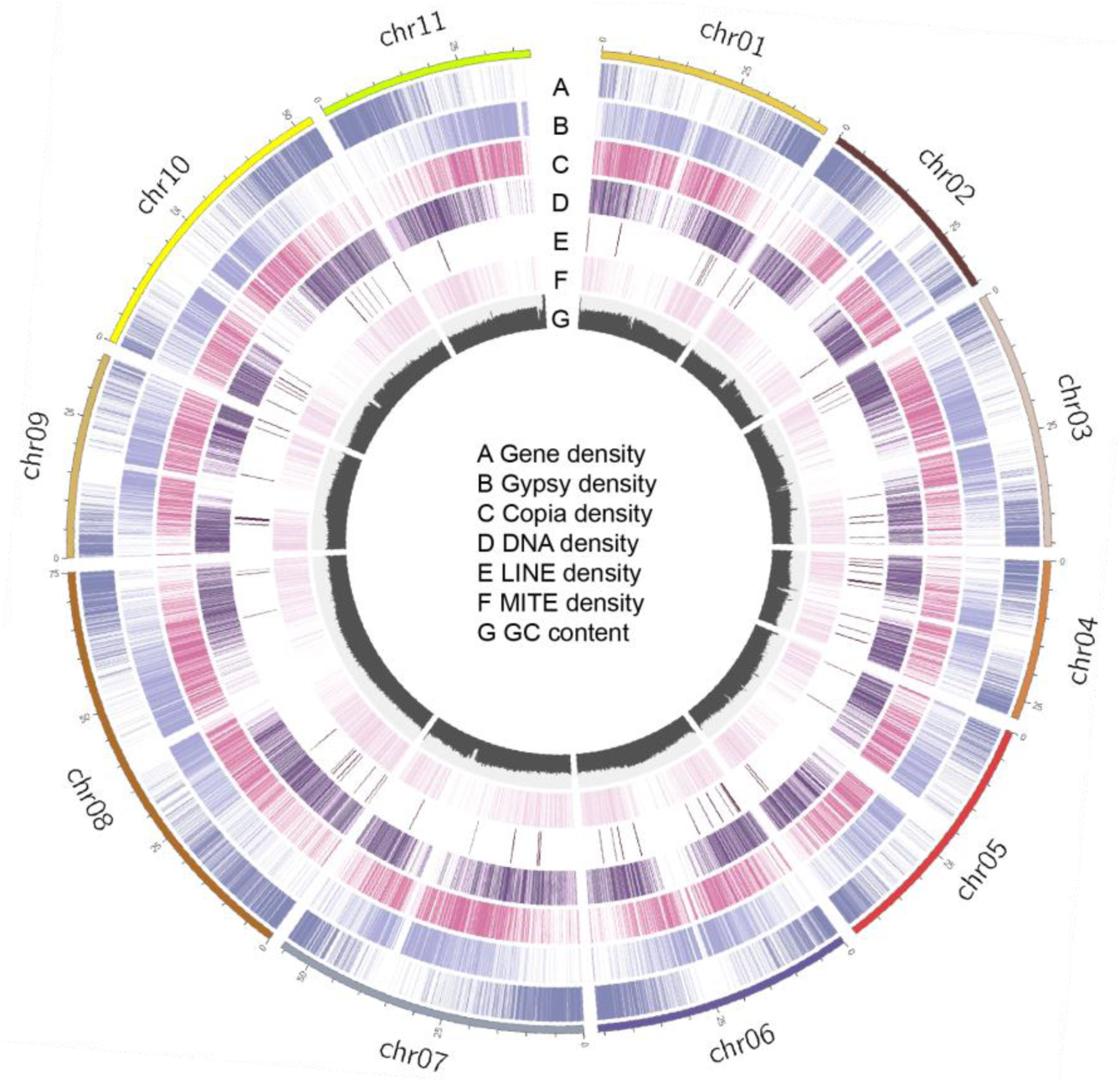
Telomere-to-telomere genomic features of mung bean. The outermost circle displays chromosome numbers and lengths. (A) Gene density; (B) Gypsy density; (C) Copia density; (D) DNA density; (E) LINE density; (F) MITE density; (G) GC content. Counts were calculated for every 100 kb and categorized into four classes based on the count numbers, with darker colors representing higher densities.

**Table 1.** Mung bean genome assembly statistics.

| Genome characteristics |  |
| --- | --- |
| Genome size (Mb) | 500 |
| Contig number | 13 |
| Number of chromosomes | 11 |
| N50 of contigs/scaffolds (Mb) | 46 |
| L50 of contigs/scaffolds (Mb) | 5 |
| BUSCO of assembled genome (%) | 98.8 |
| Number of gaps | 0 |
| Number of protein-coding genes | 28,740 |
| BUSCO of protein-coding genes (%) | 99.0 |
| Annotated protein-coding genes (%) | 98.5 |
| Repeat (%) | 49.17 |
| LTR (%) | 30.13 |

### Comprehensive genome annotation

A total of 801,958 repetitive sequences were identified, cumulatively spanning 246,179,028 bp, which constitutes nearly half (49.17%) of the genome’s total length (**Supplementary Table 2**). Of these, LTR sequences accounted for 250,426 with a combined length of 150,861,724 bp, representing 30.13% of the genome. Within the LTR category, Gypsy elements comprised 11.65%, while Copia elements constituted 8.21%. Another significant category of repetitive elements was the TIR class, occupying 8.86% of the genome.

For the comprehensive annotation of protein-coding genes in the mung bean genome, full-length transcriptome sequencing was performed. This involved pooling RNA extracted in equimolar amounts from root, stem, and leaf tissues, yielding 15G of ONT data (**Supplementary Table 1**). To enhance the robustness of gene annotation, 214 next-generation sequencing transcriptomic datasets were also compiled (**Supplementary Table 3**). In total, 28,740 protein-coding genes, 1,561 rRNAs, 586 tRNAs, and 843 ncRNAs (including miRNAs, tRNAs, snRNAs, and other small ncRNAs) were annotated in the mung bean genome (**Table 1**).

For functional annotation of these protein-coding genes, three strategies were employed. In total, 28,320 (98.54%) protein-coding genes were annotated with functional information (**Table 1, Supplementary Table 4**). The largest proportion of functionally annotated genes was found in the TrEMBL and NR databases, accounting for 97.72% and 94.61%, respectively (**Supplementary Table 4**). Additionally, functional annotations were assigned to 47.65% of genes in the GO database and 27.42% in the KEGG pathway database.

### Assessment of T2T genome completeness

We utilized the Benchmarking Universal Single-Copy Orthologs (BUSCO) assessment with the embryophyta_odb10 lineage dataset to evaluate the completeness of the mung bean genome, revealing that out of the 1,614 core genes expected, the genome showed a high level of completeness with BUSCO scores reaching 98.8% (**Table 1, Supplementary Table 5**). In a further assessment of the protein-coding genes using BUSCO, the genome displayed an even higher completeness score, with 99.0% of the core genes being fully represented (**Table 1, Supplementary Table 5**). Additionally, Merqury analysis, performed using a *k*-mer size of 19 based on HiFi reads, indicated a high level of genomic accuracy. The consensus quality values (QVs) were recorded at 64.26, and the *k*-mer completeness scores reached an impressive 99.73 (**Table 1, Supplementary Fig. 2**). Moreover, when mapping Hi-C data onto the genome, no significant assembly errors were detected, indicating a high level of structural accuracy (**Supplementary Fig. 3**). Taken together, these various assessments underscore the exceptional quality and reliability of the mung bean genome assemblies. **Recent LTR-RT expansion altering nearby gene expression**

Using SubPhaser (Jia *et al*., 2022), we identified a total of 144,186 species-specific *K*-mer sequences based on the direct homologous relationship among the chromosomes of cowpea, common bean, and mung bean genomes (**Fig. 2A**). These species-specific *K*-mer sequences are sufficient to distinguish the three genomes (**Fig. 2B-C**). The majority of these species-specific *K*-mers are distributed in the central regions of the chromosomes and map to 74,772 TE sequences, with 23,980 in mung bean, 24,777 in common bean, and 26,015 in cowpea (**Fig. 2D**). Further classification of these TEs revealed that over 99% of the species-specific amplified TEs are LTR retrotransposon (LTR-RT) sequences, consistent with previous findings in species such as wheat (Jia *et al*., 2022). In mung bean and cowpea, LTR expansion began approximately 7.9 million years ago, while in common bean, it began approximately 7 million years ago, indicating their divergence around 7-8 million years ago (**Fig. 2E**), consistent with conclusions drawn from phylogenetic analysis (Liang *et al*., 2022; Francis *et al*., 2023). The evolutionary tree of LTR-RTs shows that species-specific LTR-RTs have evolved towards different directions (**Supplementary Fig. 4**).

**Fig. 2.**
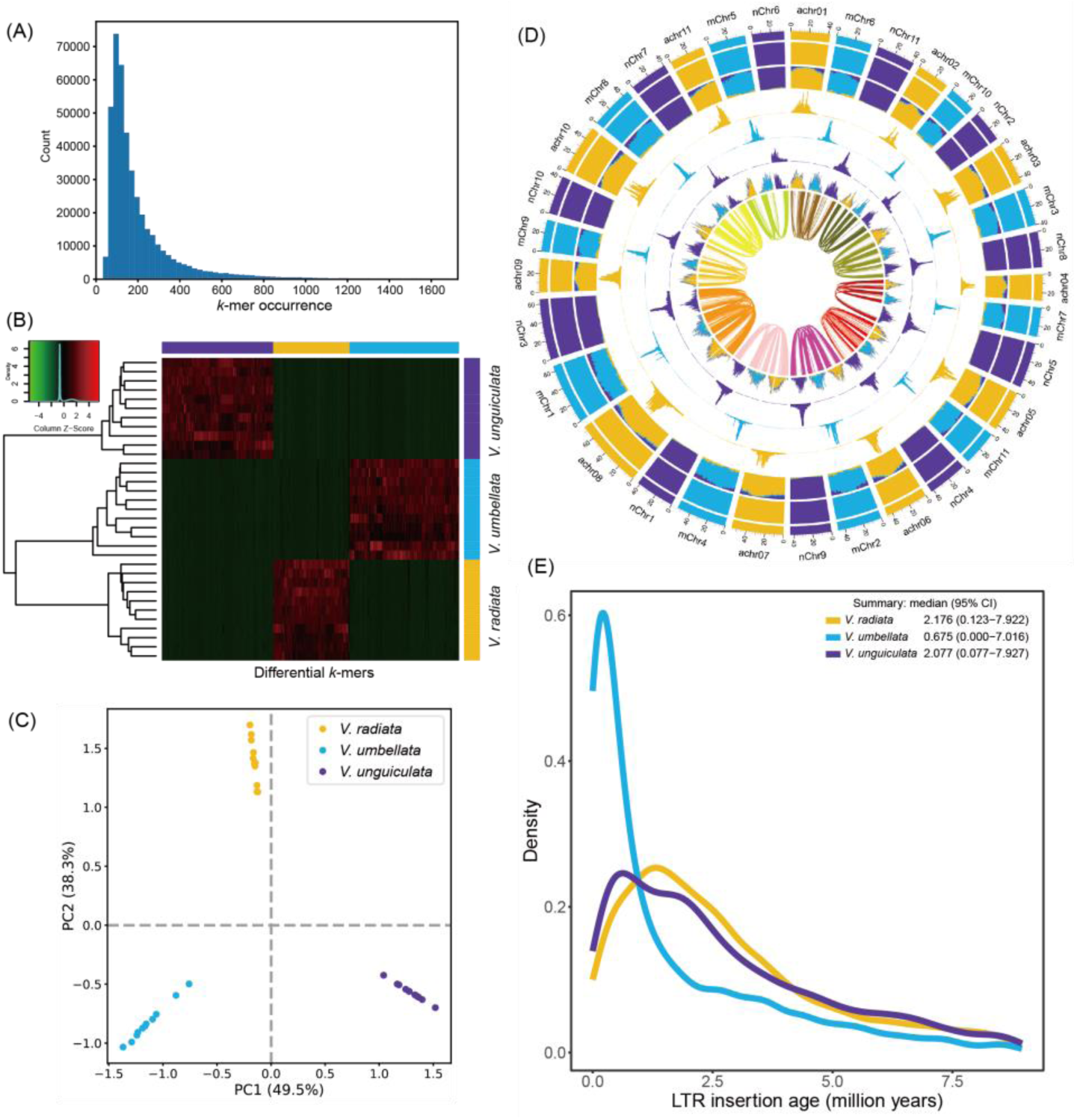
Phasing and characterization of three *Vigna* genomes. (A) The number of differential 15-mers among the homoeologous chromosomes. (B) Unsupervised hierarchical clustering is shown, with the horizontal color bar at the top indicating the specific genome of the k-mer, and the vertical color bar on the left indicating the assigned genome of the chromosome. The heat map displays the Z-scaled relative abundance of k-mers, with higher Z scores indicating greater relative abundance. (C) Principal component analysis (PCA) of differential 15-mers confirms successful phasing into three genomes, based on distinct patterns of differential k-mers and homoeologous chromosomes. (D) Chromosomal features. From the outermost to the innermost circles (1–8): (1) genome assignments based on a k-means algorithm; (2) significant enrichment of genome-specific k-mers, where the same color as the genome indicates significant enrichment, and white areas indicate no significant enrichment; (3) normalized proportion (relative) of genome-specific k-mers; (4–6) count (absolute) of each genome-specific k-mer set; (7) density of long terminal repeat retrotransposons (LTR-RTs), with consistent colors indicating significant enrichment of LTR-RTs to those genome-specific k-mers, and gray indicating nonspecific LTR-RTs; (8) homoeologous blocks. All statistics (2–7) are computed using sliding windows of 1 Mb. (E) Insertion time of genome-specific LTR-RTs, with a 95% confidence interval (CI) in the upper right corner predicting the insertion time boundaries of LTR-RTs on the genome. (b–f) Colors are consistent across genomes.

Among the 23,980 species-specific TE sequences identified in mung bean, 23,805 (99.27%) belong to the LTR class (**Fig. 3A**). Within the LTR class, Ty1/copia and Ty3/gypsy were the most abundant invasions. For instance, Ale from Ty1/copia invaded 5,353 times in mung bean, while Athila from Ty3/gypsy invaded 4,083 times (**Fig. 3A**). These specific TEs inserted into 465 genes, 678 genes within the upstream 5 kb, and 791 genes within the downstream 5 kb. Interestingly, we found that the insertion of these TEs did not specifically interfere with the function of a particular category of genes, but mainly interfered with functions such as cellular hyperosmotic salinity response, benzoate metabolic process, and positive regulation of proteolysis (**Supplementary Table 6-8**). Furthermore, to examine whether these different classes of TEs specifically inserted into certain positions, we categorized these TEs into four groups based on their abundance: Ty1_copia/Ale, Ty1_copia/Ivana, Ty3_gypsy/chromovirus/Reina, and TE/others. We found that regardless of whether they were inserted upstream, downstream, or within genes, Ty1_copia/Ale always had the highest number of insertions, followed by Ty1_copia/Ivana in upstream and downstream regions, but not as much within genes compared to Ty3_gypsy/chromovirus/Reina (**Fig 3B**). We performed enrichment analysis on the genes affected by these four classes of TEs and found distinct patterns of insertion. The Ty3_gypsy/chromovirus/Reina primarily inserted within genes associated with the cellular response to DNA damage stimulus (**Supplementary Table 9-11**). Ty1_copia/Ale mainly inserted within genes involved in vesicle-mediated transport (**Supplementary Table 12-14**). Ty1_copia/Ivana predominantly inserted within genes related to inorganic ion import across the plasma membrane (**Supplementary Table 15-17**). Notably, these TEs did not predominantly influence the function of specific gene categories in the upstream or downstream 5 kb regions (**Supplementary Table 9-17**).

**Fig. 3.**
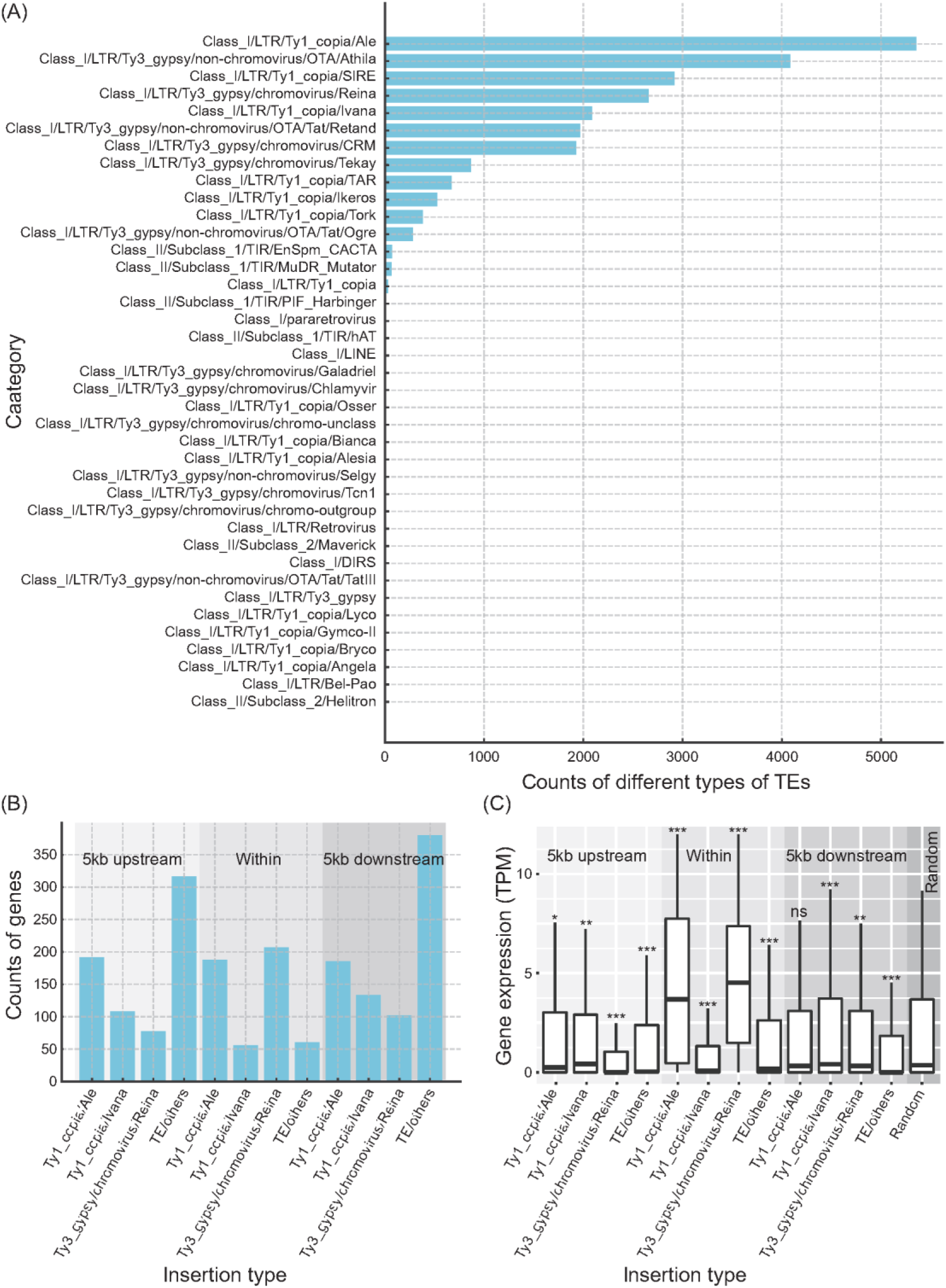
Analysis of TEs in mung bean. (A) Counts of different types of recently expanded TEs in mung bean. The x-axis represents the counts of various TEs, while the y-axis lists the different TE categories identified. (B) Distribution of different TE types within gene regions, 5kb upstream, and 5kb downstream. The x-axis represents the insertion type, and the y-axis represents the counts of genes associated with these insertions. (C) Gene expression levels (TPM) for genes with TE insertions within 5kb upstream, within gene bodies, and 5kb downstream. The box plots compare the gene expression levels with a random set of 500 genes from the genome. The significance levels indicate the statistical difference compared to the random set (\**p* < 0.05, \*\**p* < 0.01, \*\*\**p* < 0.001; Wilcoxon rank sum test).

To investigate the impact of these TEs on gene expression, we downloaded a set of transcriptome data from leaf tissues of cultivated and wild mung beans (PRJNA771920) from NCBI. Our analysis revealed that TE insertions upstream, downstream, or within genes significantly altered gene expression (**Fig. 3C**, Wilcoxon rank sum test). For instance, Ty1_copia/Ale insertions upstream of genes significantly reduced gene expression, while insertions downstream had no effect. However, insertions within the gene body significantly increased gene expression. Ty3_gypsy/chromovirus/Reina insertions both upstream and downstream of genes significantly decreased gene expression, whereas insertions within the gene body also enhanced gene expression.

### Population structure of cultivated and wild mung bean under structural variation

We collected high-depth genomic data from 36 wild and 78 cultivated mung bean samples, with an average sequencing depth of 33.5 ± 9.7× per individual (**Supplementary Table 21**). These reads were mapped to the newly assembled “Weilv-9” genome sequence, revealing an average coverage depth of 29×, ranging from 16.4× to 61.3×, covering 85%-98% of the genome (**Supplementary Table 21**). A total of 25,450,640 variant sites were identified, and after stringent filtering, 13,385,833 high-quality SNPs with a missing rate of less than 20% were retained. Further filtering for minor allele frequency (MAF) less than 0.05 resulted in 3,069,438 SNPs.

In parallel, we identified 365,831 SVs supported by at least two individuals and with a missing rate of less than 20%. This resulted in 115,398 retained SVs, with DELs being the most common type, comprising 90,180 variants. Additionally, there were 23,250 INSs, 1,290 DUPs, and 678 INVs. These SVs spanned approximately 94 Mb of the genome. Inversions had the longest average length at 17,056 bp, followed by duplications with an average length of 5,690 bp, while insertions had the shortest average length (**Supplementary Table 22**).

Based on unlinked SNPs, our population structure analysis was consistent with previous results. At *K*=2, the cultivated and wild populations split into two distinct groups (**Fig. 4A-B**), except for two individuals potentially misassigned (Lin *et al*., 2023). Similarly, SV-based analysis supported these findings. Both SNP and SV analyses indicated minimal introgression between the two populations (**Fig. 4A, C**), unlike soybean, which shows significant introgression from cultivated to wild populations (Wang *et al*., 2019; Jia *et al*., 2024). After excluding the two potentially misassigned individuals, we compared the density of SNPs and SVs, finding a strong correlation (general linear model: *t*-value = 20.84, *p* < 2e-16) (**Fig. 4D**).

**Fig. 4.**
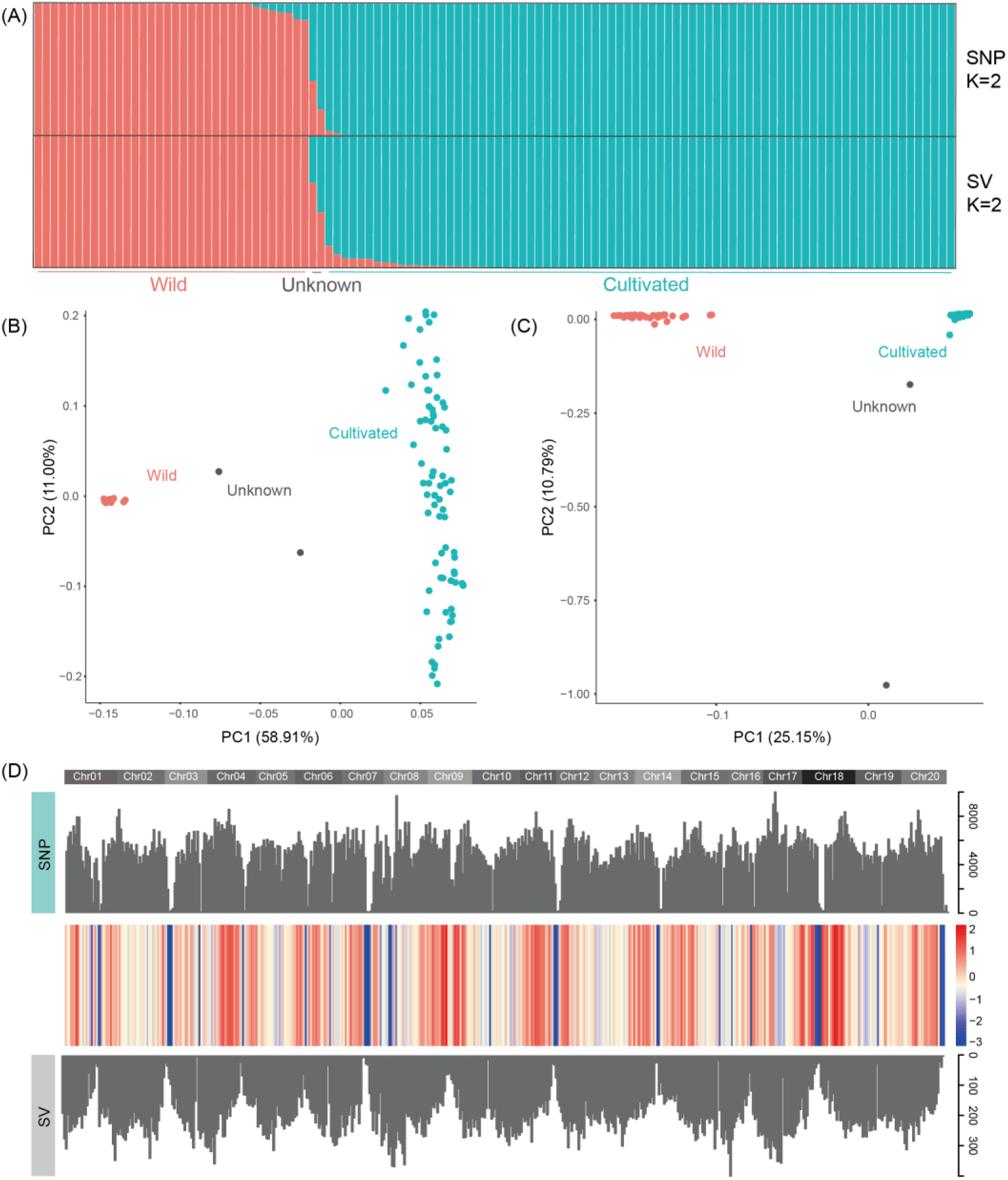
Population structure of cultivated and wild mung beans. (A) Ancestry assignment for 114 mung bean individuals based on SNPs and SVs at *K* = 2. Each bar represents an individual, with different colors indicating varying ancestry components. (B) PCA based on SNPs for cultivated and wild mung beans, with different colors representing different species/groups. (C) PCA based on SVs for cultivated and wild mung beans, with different colors representing different species/groups. (D) The residuals from a regression of SV density on SNP density (10 Mb windows), with chromosome labels above the plot. Positive residuals correspond to regions with more SVs than expected based on SNP density, while negative residuals correspond to regions with fewer SVs than expected.

### Divergence between cultivated and wild mung bean under SVs

The cultivated population harbors 1,049,411 private SNPs, which impact 21,795 genes. In contrast, the wild population contains relatively fewer private SNPs (975,936) compared to the cultivated group. Interestingly, we identified a higher number of private structural variants (SVs) in the wild population (51,811) than in the cultivated population (33,559). The site frequency spectrum (SFS) of these SNPs and SVs predominantly shows a MAF below 0.05 (**Fig. 5A-B**). Generally, low-frequency SVs are likely to be deleterious or neutral and are typically not widespread in the population due to negative selection pressure, while high-frequency SVs tend to be neutral or beneficial, allowing them to persist and propagate within the population (Pollex & Hegele, 2007; Sudmant *et al*., 2015). Consequently, we categorized these SVs into two groups based on their MAF: 0-0.05 (Minor Allele Frequency, MiAF) and 0.05-0.5 (Major Allele Frequency, MaAF). In the cultivated population, MiAF SVs predominantly affect 7,271 regions either 2kb upstream, downstream, or within genes, whereas MaAF SVs impact 5,572 genes. Similarly, in the wild population, MiAF SVs affect 8,823 genes, while MaAF SVs influence 11,730 genes. Notably, we observed that in both cultivated and wild populations, the expression levels of genes impacted by MaAF SVs were significantly lower than those affected by MiAF SVs (**Fig. 5C**).

**Fig. 5.**
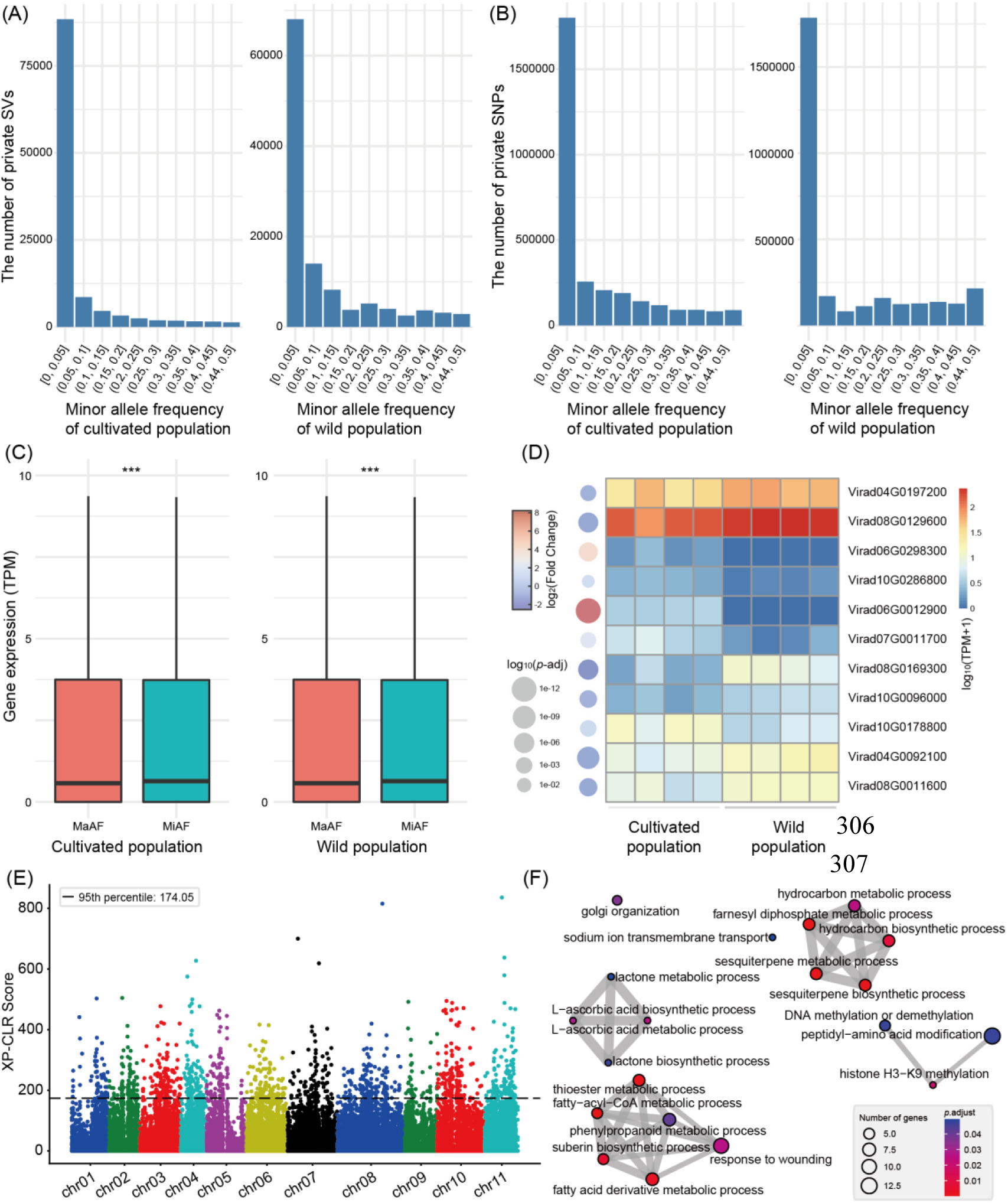
Genomic variations in cultivated and wild mung bean populations. (A) Number of private SNPs in the cultivated and wild populations. (B) Number of private SVs in the cultivated and wild populations. (C) Gene expression levels in genes affected by minor allele frequency (MiAF) and major allele frequency (MaAF) SVs in both cultivated and wild populations. Asterisks indicate significant differences (*** *p* < 0.001, Wilcoxon rank sum test). (D) The expression levels of 11 significantly differentially expressed genes (*p* < 0.05, FC > 2). (E) Genome-wide distribution of 20-kb windows based on XP-CLR scores for SNPs. (F) Enrichment analysis of biological processes among genes affected by the intersection of the top 5% of loci with the highest cross-population composite likelihood ratio (XP-CLR) scores and the structural variation dataset.

To elucidate potential genes implicated in the domestication process, we aimed to identify regions of significant chromosomal divergence between wild and cultivated samples. We estimated SNP divergences within fixed 20-kb genomic windows for this purpose. The genome-wide average fixation index (*F*_ST_) estimates were substantially higher for SNPs (0.54) than for SVs (0.44), indicative of the generally lower population frequencies observed for SVs. We identified the top 5% of windows with the highest *F*_ST_ values for both SNPs and SVs, and intersected these datasets, resulting in a set of 206 genes. This set includes genes involved in cell activation in immune response and defense response to symbionts (**Supplementary Table 23**).

Among these, 11 genes exhibited significant differential expression in leaf tissue, with 5 upregulated in cultivated varieties and 6 upregulated in wild varieties (**Fig. 5D**). Notably, two genes showed differential expression greater than fourfold. Virad06G0012900, a member of the polygalacturonase family involved in pectin metabolism, was upregulated in wild varieties. In wild environments, mung beans may require enhanced cell wall degradation capabilities to adapt to complex soil and microbial conditions, which could exert selective pressure on these genes. Conversely, cultivated mung beans, benefiting from agricultural management and fertilizer use, may experience reduced selective pressure on this gene. Virad06G0298300 contains a C2 domain, akin to synaptotagmins, which plays a crucial role in signal transduction and membrane fusion. Wild mung beans might need rapid environmental signal responses to cope with various stresses, whereas cultivated mung beans in more stable environments may not require such rapid signal transduction.

### Selective pressures in the domestication of mung beans under SVs

A cross-population composite likelihood ratio (XP-CLR) test was utilized to identify species-specific selective sweeps in mung beans based on the SNP dataset. The analysis prioritized the top 5% of 20-kb windows based on the highest XP-CLR scores, resulting in 1,074 regions (**Fig. 5E**). These regions were further intersected with SVs, yielding 382 SVs involving 350 genes. These genes were significantly enriched in fatty-acyl-CoA metabolic process, farnesyl diphosphate metabolic process, fatty acid derivative metabolic process, sesquiterpene biosynthetic and metabolic processes, and phenylpropanoid metabolic process (**Fig. 5F**). This suggests that fatty acid synthesis or degradation has undergone selective pressure. Additionally, processes such as suberin biosynthetic process and phenylpropanoid metabolic process were also strongly selected, indicating their critical roles in the domestication of mung beans from prostrate to erect growth forms.

Suberin, a biopolymer composed of phenolic and fatty acid derivatives, is primarily deposited in the root bark, seed coats, and tuber skins of plants. It plays a vital role in preventing water loss and pathogen invasion in the plant cell wall (Nomberg *et al*., 2022). In cultivated mung beans, the selective expression of genes related to suberin biosynthesis may have contributed to increased physical strength, aiding the transition from prostrate growth to erect growth forms. Phenylpropanoid metabolic products, including lignin and flavonoids, are crucial for plant structure and defense. Lignin provides physical support and stress resistance in the cell wall, while flavonoids offer antioxidant and antimicrobial properties (Xin & Herburger, 2021). In mung beans, the selective pressure on genes involved in the phenylpropanoid metabolic process may have enhanced structural stability and disease resistance, promoting the transition from prostrate to erect growth.

## Discussion

Since the first chromosome-level genome assembly of mung bean was published in 2014 (Kang *et al*., 2014), three versions have been released to date (Kang *et al*., 2014; Ha *et al*., 2021; Liu *et al*., 2022). However, these assemblies still contain numerous gaps and unanchored regions, which can hinder the identification of functional genes and impede progress in molecular breeding. Advances in sequencing and assembly technologies, particularly the advent of PacBio HiFi, ultra-long ONT, Hi-C sequencing technologies, and the hifiasm assembler (Cheng *et al*., 2021), have enabled the creation of gap-free, T2T genomes (Li & Durbin, 2024). In this study, we successfully assembled the first T2T, gap-free genome of mung bean using PacBio HiFi, ultra-long ONT, and Hi-C sequencing technologies, along with the upgraded hifiasm software. This genome assembly is also the first T2T genome for a *Vigna* species. The comprehensive assembly includes 11 chromosomes and the complete chloroplast and mitochondrial genomes, spanning a total length of 500 Mb with an impressive contig/scaffold N50 of 45 Mb. The high quality of the assembly is underscored by the BUSCO scores, indicating a completeness of 98.8% for the genome. The structural accuracy of the assembly is validated by the identification of telomeric repeats and the Hi-C interaction maps.

In addition to genome assembly, accurate gene annotation is crucial. In *V. stipulacea*, 30,963 protein-coding genes have been annotated (Takahashi *et al*., 2023), while in ricebean (*V. umbellata*), the number ranges from 31,276 to 37,489 (Kaul *et al*., 2022; Francis *et al*., 2023). For Black gram (*V. mungo*), annotations range from 28,881 to 42,115 protein-coding genes (Jegadeesan *et al*., 2021; Ambreen *et al*., 2022). Previous mung bean genome assemblies have annotated 22,427(Kang *et al*., 2014), 30,958 (Ha *et al*., 2021), and 40,125 (Liu *et al*., 2022) protein-coding genes. The coverage and continuity of the genome, annotation algorithms and tools, prediction models, and training datasets all impact the accuracy of protein-coding gene annotations (Abbas *et al*., 2024). To enhance annotation accuracy, we downloaded 213 next-generation RNA-seq datasets from NCBI and generated 15Gb of full-length transcriptome data. Additionally, we incorporated 237,544 non-redundant protein sequences from species such as *V. unguiculata*, *V. angularis*, *V. radiata*, *V. subterranea*, *V. umbellata*, *Glycine max*, *Arachis hypogaea*, *Cicer arietinum*, *Medicago truncatula*, and *Arabidopsis thaliana* as homologous protein evidence for gene annotation. Ultimately, the comprehensive annotation of 28,740 protein-coding genes and various non-coding RNAs enhances our understanding of the mung bean genome’s functional landscape. BUSCO assessment revealed that 99.0% of our annotated protein-coding genes were complete, indicating high annotation accuracy. Additionally, using multiple databases, we achieved high-quality functional annotations for 98.54% of the genes.

The detailed gene annotation lays the groundwork for functional genomics studies aimed at elucidating the roles of specific genes and their regulatory networks. For instance, the identification of genes involved in stress responses, metabolism, and signaling pathways can guide targeted breeding efforts to develop mung bean varieties with improved stress tolerance, disease resistance, and nutritional quality. Moreover, the discovery of TEs such as Ty1_copia/Ale and Ty3_gypsy/chromovirus/Reina within gene regions and their impact on gene expression suggests that TEs play a significant role in regulating gene function and genome evolution. Understanding the mechanisms by which TEs influence gene expression and contribute to phenotypic diversity can provide new strategies for crop improvement (Hassan *et al*., 2023; He *et al*., 2024).

High-throughput sequencing technologies, especially those targeting SNPs, have become essential tools for studying the domestication process. Large-scale SNP analysis enables the identification of genes selected during domestication and the loci potentially lost due to artificial selection. This provides valuable insights into the genetic basis of crop domestication. However, our understanding of SVs in this context remains relatively limited. SVs, due to their complexity and size, have been significantly underestimated in their potential impact on genetic improvement (Scott *et al*., 2021). SVs can cause significant changes in gene expression, affecting plant traits and adaptability. For example, during tomato domestication, certain SVs have impacted key genes regulating fruit development, leading to the evolution of diverse large-fruited varieties from small-fruited wild types (Alonge *et al*., 2020; Jobson & Roberts, 2022). Similarly, in cotton improvement studies, GWAS based on SVs have identified SVs associated with fiber quality and yield (He *et al*., 2021). Interestingly, some related genes were not identified using SNP-based GWAS, indicating that SVs and SNPs reveal different genetic information. Additionally, a tomato GWAS based on 17 flavor volatiles and 249 metabolites found only 5.2% overlap in quantitative trait loci between SVs and SNPs (Li *et al*., 2023). Another study on millet also found that SV-based GWAS improved the association efficiency for certain traits, with approximately 36.9% of SVs located more than 50kb away from SNPs (He *et al*., 2023).

In our study, we found that the cultivated mung bean population had a significantly lower number of SVs and private SVs compared to the wild population, despite the fact that the number of wild individuals we used was half that of the cultivated individuals. This result, consistent with SNP-based findings, further supports the impact of selective pressures on genetic diversity during domestication. Furthermore, our analysis categorized these SVs into two groups based on their MAF: 0-0.05 (MiAF) and 0.05-0.5 (MaAF). In the cultivated population, MiAF SVs predominantly affect 7,271 regions within 2kb upstream or downstream of genes, whereas MaAF SVs impact 5,572 genes. Similarly, in the wild population, MiAF SVs affect 8,823 genes while MaAF SVs influence 11,730 genes. Notably, the expression levels of genes impacted by MaAF SVs were significantly lower than those affected by MiAF SVs in both populations, suggesting a potential regulatory role of these variants in gene expression. These findings indicate that the conservation of wild mung bean populations is critical for maintaining the genetic diversity necessary for future breeding and adaptation efforts. Wild populations serve as a genetic reservoir that can be tapped into to enhance the genetic robustness of cultivated varieties. Protecting these natural habitats and preventing genetic erosion through conservation strategies will ensure the availability of diverse genetic material for sustainable agriculture.

To elucidate potential genes implicated in the domestication process, we identified regions of significant chromosomal divergence between wild and cultivated samples. Using *F*_ST_ estimates within fixed 20-kb genomic windows, we found that the genome-wide average *F*_ST_ was substantially higher for SNPs (0.54) than for SVs (0.44). This suggests a generally lower population frequency of SVs compared to SNPs. By intersecting the top 5% of windows with the highest *F*_ST_ for SNPs with the SV dataset, we identified 206 genes involved in immune-related functions, some of which exhibited significant differential expression between the two populations. Future research should focus on characterizing the specific genomic regions affected by domestication and the functional implications of these changes. Investigating the interaction between SNPs and SVs in shaping phenotypic diversity will provide deeper insights into the genetic mechanisms underlying adaptation and domestication. Additionally, studies exploring the epigenetic modifications associated with structural variants could reveal further layers of regulation that contribute to the observed differences in gene expression.

The XP-CLR analysis identified 1,074 regions in mung beans, with 382 SVs involving 350 genes, significantly enriched in pathways such as fatty-acyl-CoA metabolic process, farnesyl diphosphate metabolic process, and phenylpropanoid metabolic process. This suggests selective pressures on fatty acid metabolism and secondary metabolite production, essential for plant defense and structural integrity. The strong selection on suberin and phenylpropanoid pathways indicates their role in the transition from prostrate to erect growth forms, enhancing physical strength and pathogen resistance. Studies in maize, poplar, and rice have shown similar impacts of lignin biosynthesis gene mutations on plant mechanical properties and disease resistance, supporting the importance of these pathways in mung bean domestication (Xiong *et al*., 2019; Qin *et al*., 2020; Ma, 2024; Wang *et al*., 2024).

## Methods

A new variety of mung bean, Weilv-9, developed by the Agricultural Institute of Weifang, Shandong Province, was germinated in the laboratory. Fresh leaves were harvested for DNA extraction using a modified CTAB method as previously described (Doyle & Doyle, 1987).

Three sequencing approaches were employed to analyze the genomic DNA. Initially, DNA integrity was assessed using Femto Pulse, followed by library preparation with the Pacific Biosciences SMRTbell Express Template Prep Kit 2.0, and sequencing was conducted on the PacBio Sequel II platform. Additionally, DNA fragments that passed quality control were utilized for library construction using the Oxford Nanopore SQK-LSK109 kit and were sequenced on the PromethION sequencer (Oxford Nanopore Technologies, Oxford, UK) using R9.4 flow cells. Finally, Hi-C libraries were generated following the protocol by Wang et al. (Wang *et al*., 2015) and sequenced using the DNBSEQ platform.

### *De novo* genome assembly

The genome was assembled utilizing a hybrid strategy that incorporated PacBio HiFi, ONT, and Hi-C sequencing reads. Initial contigs were constructed using hifiasm v0.19.8-r602 (Cheng *et al*., 2021). Chromosomal-scale scaffolding commenced with the alignment of Hi-C reads to the primary assembly using Juicer v1.6 (Durand *et al*., 2016). This was followed by clustering with 3D-DNA v180922 (Dudchenko *et al*., 2017) and manual refinement of potential mis-assemblies using Juicebox v1.11.08 (Robinson *et al*., 2018). The process resulted in chromosome-scale scaffolds and several unplaced contigs, with gaps of 100bp. These gaps were subsequently closed using LR_Gapcloser (Xu *et al*., 2019), leveraging Nanopore reads. Telomere sequences were extended by realigning HiFi reads to the chromosomal ends, thus extending the scaffolds. Organelle genomes were independently assembled using GetOrganelle (Jin *et al*., 2020). The final genome assembly was polished with a single round of NextPolish v1.3.1 (Hu *et al*., 2020) using short reads. To eliminate redundancies and contaminants, unplaced contigs were aligned to the chromosomal and organelle assemblies using redundans v0.13c (Pryszcz & Gabaldon, 2016) to identify and subsequently remove haplotigs, organellar fragments, and rDNA fragments through manual inspection. This integrated approach of long-read sequencing and Hi-C scaffolding culminated in a robust, high-quality, chromosome-scale reference genome.

### Repeat identification

TEs were *de novo* identified from the genome using the EDTA v1.9.9 (Ou *et al*., 2019). The analysis was conducted under sensitive settings (––sensitive 1 ––anno 1) to develop a comprehensive TE library. This library served as a reference for RepeatMasker (Chen, 2004), which facilitated the identification and annotation of repetitive elements within the genome. Using the consensus sequences and profile hidden Markov models from the EDTA-generated library, RepeatMasker executed sequence alignments to classify and delineate genomic repeats, including low-complexity DNA, satellite sequences, simple repeats, and transposable elements.

### Transcriptome data sequencing and collection

To elucidate the transcriptomic profile of wild mung bean, nanopore sequencing was employed on a pooled sample consisting of roots, stems, and leaves, generating 15G of full-length cDNA reads. Additionally, this analysis was enhanced by incorporating 214 high-quality short-read RNA-seq datasets sourced from the NCBI Sequence Read Archive.

### Gene prediction and function annotation

The genome was annotated by leveraging evidence from homologous proteins, transcript data, and *ab initio* predictions. Homologous protein evidence was gathered by searching against a collection of 237,544 non-redundant protein sequences from ten plant species, including *Vigna* and *Glycine* genera, among others. Transcript evidence was procured through alignment of both short and long RNA-seq reads to the genome, using Hisat2 v2.1.0 (Kim *et al*., 2015) for short reads and minimap2 v2.24-r1122 (Li, 2018) for long reads, followed by assembly with StringTie v1.3.5 (Pertea *et al*., 2015).

Gene structures were annotated using PASA v2.4.1 (Haas *et al*., 2003), which utilized transcript evidence. Full-length genes were identified by comparing them to reference proteins. AUGUSTUS v3.4.0 (Stanke *et al*., 2008) was then trained and optimized for gene prediction using the identified full-length gene set. The MAKER v2.31.9 (Cantarel *et al*., 2008) annotation pipeline integrated *ab initio* predictions, transcript, and protein evidence. EvidenceModeler (EVM) v1.1.1 (Haas *et al*., 2008) was employed to integrate and refine the MAKER and PASA annotations, with TEsorter (Zhang *et al*., 2022) used to mask TE protein domains. Abnormal gene models and those under 50 amino acids were excluded.

Non-coding RNA (ncRNA) annotations were performed using tRNAScan-SE v1.3.1 (Lowe & Eddy, 1997), barrnap (https://github.com/tseemann/barrnap), and RfamScan (Kalvari *et al*., 2018). Functional annotations of protein-coding genes were executed using three complementary approaches. First, alignment to the eggNOG v5.0 ortholog database using eggNOG-mapper v2.0.1 (Cantalapiedra *et al*., 2021) provided GO and KEGG annotations. Secondly, similarity searches against protein databases such as SwissProt, TrEMBL, and others were conducted with diamond v0.9.24 (Buchfink *et al*., 2015), identifying best-matching orthologs based on sequence identity and E-value criteria. Finally, domain architecture analysis was carried out by scanning against Hidden Markov Models in the InterPro database using InterProScan v5.27-66.0 (Jones *et al*., 2014). These homology-based methods collectively facilitated a comprehensive functional annotation of the protein-coding genes, linking them to known sequences and conserved domains.

### Assessment of genome completeness and protein-coding genes

The genome assembly’s quality was assessed through three distinct methodologies. Initially, the completeness of the genome sequences and protein-coding gene sets was evaluated using BUSCO v2.0.1 (Simão *et al*., 2015), which quantifies the presence of conserved core genes across the assembly. Subsequently, the assembly’s fidelity was gauged using Merqury (Rhie *et al*., 2020), which employs a k-mer size of 19 to evaluate genome accuracy. Lastly, to detect potential assembly errors, Hi-C data was mapped onto the assembled genome using Juicer, providing insights into the structural integrity of the genome.

### Genome comparison and species-specific sequence identification

SubPhaser v1.0.5 (Jia *et al*., 2022) was employed to identify differential repeats across the three *Vigna* genomic assemblies. Bedtools (Quinlan, 2014) was used to ascertain the intersections between functional genes and specific repetitive sequences.

### Transcriptome data analysis

Fastp v0.12.4 (Chen *et al*., 2018) is utilized to eliminate short reads (less than 60 bp), potential adapter sequences, and low-quality reads from the sequencing data. Following this preprocessing, gene expression levels are quantified in transcripts per million (TPM) using salmon v1.6.0 (Patro *et al*., 2017), which incorporates advanced options such as ––validateMappings and ––numBootstraps 100 to ensure accurate mapping and estimation of expression variability. The processed data is then further analyzed with tximport v1.22.0 (Soneson *et al*., 2015) for downstream bioinformatics evaluations.

### Collection and SNP calling of whole-genome resequencing data

A total of 114 mung bean accessions, including 36 wild and 78 cultivated varieties, were downloaded from NCBI’s project PRJNA838242 (Lin *et al*., 2023). Paired-end reads were processed using fastp v0.23.2 to trim adapters, remove low-quality bases, and discard short reads (less than 60 bp). These filtered reads were then aligned to the “Weilv-9” reference genome using BWA-MEM (Li, 2013). SNP calling was carried out with Freebayes (Garrison & Marth, 2012). To minimize bias in SNP calling and ensure only high-quality SNPs were retained, we implemented the following filtering steps: (1) SNPs with a quality score below 20 were removed; (2) only bi-allelic SNPs were retained; (3) depth parameters were set to ––minDP 5 and ––maxDP 500; (4) SNPs with a missing rate greater than 20% were removed; (5) all SNPs with a MAF below 0.05 were also excluded.

### SV discovery

In Dysgu v1.6.3 (Cleal & Baird, 2022), each BAM file was processed using the dysgu run command to identify SVs, retaining only those flagged as “PASS”. The filtered VCF files from all samples were then merged using the dysgu merge command. Subsequently, a second round of SV calling was performed on the merged VCF using dysgu run ––sites, with only variants meeting the established filtering criteria retained. The final consolidated VCF file was generated by merging the filtered VCF files from all individuals once again using dysgu merge. We then removed variants with a missing rate greater than 20%, those with a minimum allele count less than 2, and SVs larger than 1 Mb.

### Examining SNP and SV density patterns

To investigate the correlations between SNP and SV densities throughout the mung bean genome, we utilized tabix v1.9 (Li, 2011) to quantify variants within 1 Mb intervals across all 20 chromosomes. We conducted correlation analyses using the glm() function, specifically focusing on the residuals retained from these binned variant densities. The overlap among different genomic regions was assessed using the bedtools intersect function.

### Population structure and divergence

We utilized ADMIXTURE v1.3.0 (Alexander *et al*., 2009) for population structure analysis and GCTA v1.94.1 (Yang *et al*., 2011) for PCA. Visualization was conducted through clumak (Kopelman *et al*., 2015) and ggplot2 v3.4.2 (Villanueva & Chen, 2019). We utilized vcftools v0.1.16 (Danecek *et al*., 2011) to calculate *F*_ST_, allele frequencies and private alleles.

### Putative selective sweeps

Comprehensive genome analysis was conducted employing XP-CLR (Chen *et al*., 2010), a technique predicated on the probability modeling of multilocus allele frequency divergence across two distinct populations. Following phasing with Beagle v5.4 (Browning & Browning, 2007), the XP-CLR program was executed for each chromosome with specified parameters: ––ld 0.95 ––phased.

### GO enrichment

GO enrichment was executed via R’s clusterProfiler v4.2.2 package (Wu *et al*., 2021). Key steps included initial GO term identification using “enrichGO”. Redundant terms were minimized using the “simplify” function, followed by inter-term similarity assessment with the “pairwise_termsim” method employing the Jaccard coefficient. Visualization of the intricate GO term network was achieved with enrichplot v1.14.2 (Yu, 2021).

## Data Availability statement

The whole genome sequence data reported in this paper have been deposited in the Genome Sequence Archive, under accession number PRJCA021300. The genome assembly and annotation data reported in this paper have been deposited in the Genome Warehouse in National Genomics Data Center, Beijing Institute of Genomics, Chinese Academy of Sciences / China National Center for Bioinformation, under accession number GWHEQVC00000000 that is publicly accessible at https://ngdc.cncb.ac.cn/gwh.

## Contributions

KHJ and NNL conceived and designed the study; KHJ, GL, LW, ML, ZWW, RZL, LLL, KX, YYY, RMT, XC, YJS, XYZ, and FJS collected materials and analyzed the data; KHJ, LW, and LLL prepared figures and tables; KHJ, LW, and GL wrote and revised the manuscript; all authors approved the final manuscript.

## Reference

1. Abbas Q, Wilhelm M, Kuster B, Poppenberger B, Frishman D. 2024. Exploring crop genomes: assembly features, gene prediction accuracy, and implications for proteomics studies. BMC Genomics 25: 619.

2. Alexander DH, Novembre J, Lange K. 2009. Fast model-based estimation of ancestry in unrelated individuals. Genome Research 19: 1655–1664.

3. Alonge M, Wang X, Benoit M, Soyk S, Pereira L, Zhang L, Suresh H, Ramakrishnan S, Maumus F, Ciren D, et al. 2020. Major impacts of widespread structural variation on gene expression and crop improvement in tomato. Cell 182: 145–161 e123.

4. Ambreen H, Oraon PK, Wahlang DR, Satyawada RR, Katiyar-Agarwal S, Agarwal M, Jagannath A, Kumar A, Budhwar R, Shukla RN, et al. 2022. Long-read-based draft genome sequence of Indian black gram IPU-94-1 ‘Uttara’: Insights into disease resistance and seed storage protein genes. The plant genome 15: e20234.

5. Browning SR, Browning BL. 2007. Rapid and accurate haplotype phasing and missing-data inference for whole-genome association studies by use of localized haplotype clustering. The American Journal of Human Genetics 81: 1084–1097.

6. Buchfink B, Xie C, Huson DH. 2015. Fast and sensitive protein alignment using DIAMOND. Nature Methods 12: 59–60.

7. Cantalapiedra CP, Hernández-Plaza A, Letunic I, Bork P, Huerta-Cepas J. 2021. eggNOG-mapper v2: functional annotation, orthology assignments, and domain prediction at the metagenomic scale. Molecular Biology and Evolution 38: 5825–5829.

8. Cantarel BL, Korf I, Robb SM, Parra G, Ross E, Moore B, Holt C, Sanchez Alvarado A, Yandell M. 2008. MAKER: an easy-to-use annotation pipeline designed for emerging model organism genomes. Genome Research 18: 188–196.

9. Chaisson MJP, Sanders AD, Zhao X, Malhotra A, Porubsky D, Rausch T, Gardner EJ, Rodriguez OL, Guo L, Collins RL, et al. 2019. Multi-platform discovery of haplotype-resolved structural variation in human genomes. Nature Communications 10: 1784.

10. Chen H, Patterson N, Reich D. 2010. Population differentiation as a test for selective sweeps. Genome Research 20: 393–402.

11. Chen N. 2004. Using RepeatMasker to identify repetitive elements in genomic sequences. Curr Protoc Bioinformatics Chapter 4: Unit 4 10.

12. Chen S, Zhou Y, Chen Y, Gu J. 2018. fastp: an ultra-fast all-in-one FASTQ preprocessor. Bioinformatics 34: i884–i890.

13. Cheng H, Concepcion GT, Feng X, Zhang H, Li H. 2021. Haplotype-resolved de novo assembly using phased assembly graphs with hifiasm. Nature Methods 18: 170–175.

14. Cleal K, Baird DM. 2022. Dysgu: efficient structural variant calling using short or long reads. Nucleic Acids Research 50: e53.

15. Cumer T, Boyer F, Pompanon F. 2021. Genome-wide detection of structural variations reveals new regions associated with domestication in small ruminants. Genome biology and evolution 13: evab165.

16. Danecek P, Auton A, Abecasis G, Albers CA, Banks E, DePristo MA, Handsaker RE, Lunter G, Marth GT, Sherry ST, et al. 2011. The variant call format and VCFtools. Bioinformatics 27: 2156–2158.

17. Doyle JJ, Doyle JL. 1987. A rapid DNA isolation procedure for small quantities of fresh leaf tissue. Phytochemical Bulletin 19: 11–15.

18. Dudchenko O, Batra SS, Omer AD, Nyquist SK, Hoeger M, Durand NC, Shamim MS, Machol I, Lander ES, Aiden AP, et al. 2017. De novo assembly of the Aedes aegypti genome using Hi-C yields chromosome-length scaffolds. Science 356: 92–95.

19. Durand NC, Shamim MS, Machol I, Rao SS, Huntley MH, Lander ES, Aiden EL. 2016. Juicer provides a one-click system for analyzing loop-resolution Hi-C experiments. Cell Syst 3: 95–98.

20. Francis A, Singh NP, Singh M, Sharma P, Gayacharan, Kumar D, Basu U, Bajaj D, Varshney N, Joshi DC, et al. 2023. The ricebean genome provides insight into *Vigna* genome evolution and facilitates genetic enhancement. Plant Biotechnology Journal 21: 1522–1524.

21. Garrison E, Marth G. 2012. Haplotype-based variant detection from short-read sequencing. arXiv preprint arXiv 1207: 3907.

22. Gaut BS, Seymour DK, Liu Q, Zhou Y. 2018. Demography and its effects on genomic variation in crop domestication. Nature Plants 4: 512–520.

23. Ha J, Lee S-H. 2019. Mung bean (*Vigna radiata* (L.) R. Wilczek) breeding. Advances in Plant Breeding Strategies: Legumes: Volume 7 7: 371–407.

24. Ha J, Satyawan D, Jeong H, Lee E, Cho KH, Kim MY, Lee SH. 2021. A near-complete genome sequence of mungbean (*Vigna radiata* L.) provides key insights into the modern breeding program. The plant genome 14: e20121.

25. Haas BJ, Delcher AL, Mount SM, Wortman JR, Smith RK, Jr., Hannick LI, Maiti R, Ronning CM, Rusch DB, Town CD, et al. 2003. Improving the *Arabidopsis* genome annotation using maximal transcript alignment assemblies. Nucleic Acids Research 31: 5654–5666.

26. Haas BJ, Salzberg SL, Zhu W, Pertea M, Allen JE, Orvis J, White O, Buell CR, Wortman JR. 2008. Automated eukaryotic gene structure annotation using EVidenceModeler and the program to assemble spliced alignments. Genome Biology 9: R7.

27. Hassan AH, Mokhtar MM, El Allali A. 2023. Transposable elements: multifunctional players in the plant genome. Frontiers in Plant Science 14: 1330127.

28. He Q, Tang S, Zhi H, Chen J, Zhang J, Liang H, Alam O, Li H, Zhang H, Xing L, et al. 2023. A graph-based genome and pan-genome variation of the model plant *Setaria*. Nature Genetics 55: 1232–1242.

29. He S, Sun G, Geng X, Gong W, Dai P, Jia Y, Shi W, Pan Z, Wang J, Wang L, et al. 2021. The genomic basis of geographic differentiation and fiber improvement in cultivated cotton. Nature Genetics 53: 916–924.

30. He X, Qi Z, Liu Z, Chang X, Zhang X, Li J, Wang M. 2024. Pangenome analysis reveals transposon-driven genome evolution in cotton. BMC Biology 22: 92.

31. Hitte C. 2019. Toward the identification and role of structural variations during dog domestication. Natl Sci Rev 6: 123–124.

32. Hou D, Yousaf L, Xue Y, Hu J, Wu J, Hu X, Feng N, Shen Q. 2019. Mung bean (*Vigna radiata* L.): bioactive polyphenols, polysaccharides, peptides, and health benefits. Nutrients 11: 1238.

33. Hu J, Fan J, Sun Z, Liu S. 2020. NextPolish: a fast and efficient genome polishing tool for long-read assembly. Bioinformatics 36: 2253–2255.

34. Ignacimuthu S, Babu C. 1987. *Vigna radiata* var. *sublobata* (Fabaceae): Economically useful wild relative of urd and mung beans. Economic Botany 41: 418–422.

35. Jegadeesan S, Raizada A, Dhanasekar P, Suprasanna P. 2021. Draft genome sequence of the pulse crop blackgram [*Vigna mungo* (L.) Hepper] reveals potential R-genes. Scientific Reports 11: 11247.

36. Jia KH, Wang ZX, Wang L, Li GY, Zhang W, Wang XL, Xu FJ, Jiao SQ, Zhou SS, Liu H, et al. 2022. SubPhaser: a robust allopolyploid subgenome phasing method based on subgenome-specific *k*-mers. New Phytologist 235: 801–809.

37. Jia KH, Zhang X, Li LL, Shi TL, Liu D, Yang Y, Cong Y, Li R, Pu Y, Gong Y, et al. 2024. Telomere-to-telomere genome assemblies of cultivated and wild soybean provide insights into evolution and domestication under structural variation. Plant Communications 10: 100919.

38. Jin JJ, Yu WB, Yang JB, Song Y, dePamphilis CW, Yi TS, Li DZ. 2020. GetOrganelle: a fast and versatile toolkit for accurate de novo assembly of organelle genomes. Genome Biology 21: 241.

39. Jobson E, Roberts R. 2022. Genomic structural variation in tomato and its role in plant immunity. Mol Hortic 2: 7.

40. Jones P, Binns D, Chang HY, Fraser M, Li W, McAnulla C, McWilliam H, Maslen J, Mitchell A, Nuka G, et al. 2014. InterProScan 5: genome-scale protein function classification. Bioinformatics 30: 1236–1240.

41. Kalvari I, Argasinska J, Quinones-Olvera N, Nawrocki EP, Rivas E, Eddy SR, Bateman A, Finn RD, Petrov AI. 2018. Rfam 13.0: shifting to a genome-centric resource for non-coding RNA families. Nucleic Acids Research 46: D335–D342.

42. Kang YJ, Kim SK, Kim MY, Lestari P, Kim KH, Ha BK, Jun TH, Hwang WJ, Lee T, Lee J, et al. 2014. Genome sequence of mungbean and insights into evolution within *Vigna* species. Nature Communications 5: 5443.

43. Kaul T, Easwaran M, Thangaraj A, Meyyazhagan A, Nehra M, Raman NM, Verma R, Sony SK, Abdel KF, Bharti J, et al. 2022. De novo genome assembly of rice bean (Vigna umbellata) – a nominated nutritionally rich future crop reveals novel insights into flowering potential, habit, and palatability centric – traits for efficient domestication. Frontiers in Plant Science 13: 739654.

44. Kim D, Langmead B, Salzberg SL. 2015. HISAT: a fast spliced aligner with low memory requirements. Nature Methods 12: 357–360.

45. Kopelman NM, Mayzel J, Jakobsson M, Rosenberg NA, Mayrose I. 2015. Clumpak: a program for identifying clustering modes and packaging population structure inferences across *K*. Molecular Ecology Resources 15: 1179–1191.

46. Li H. 2011. Tabix: fast retrieval of sequence features from generic TAB-delimited files. Bioinformatics 27: 718–719.

47. Li H. 2013. Aligning sequence reads, clone sequences and assembly contigs with BWA-MEM. arXiv preprint arXiv 1303: 3997.

48. Li H. 2018. Minimap2: pairwise alignment for nucleotide sequences. Bioinformatics 34: 3094–3100.

49. Li H, Durbin R. 2024. Genome assembly in the telomere-to-telomere era. Nature Reviews: Genetics 1: 1–13.

50. Li N, He Q, Wang J, Wang B, Zhao J, Huang S, Yang T, Tang Y, Yang S, Aisimutuola P, et al. 2023. Super-pangenome analyses highlight genomic diversity and structural variation across wild and cultivated tomato species. Nature Genetics 55: 852–860.

51. Liang L, Zhang J, Xiao J, Li X, Xie Y, Tan H, Song X, Zhu L, Xue X, Xu L, et al. 2022. Genome and pan-genome assembly of asparagus bean (*Vigna unguiculata* ssp. *sesquipedialis*) reveal the genetic basis of cold adaptation. Frontiers in Plant Science 13: 1059804.

52. Lin YP, Chen HW, Yeh PM, Anand SS, Lin J, Li J, Noble T, Nair R, Schafleitner R, Samsononova M, et al. 2023. Demographic history and distinct selection signatures of two domestication genes in mungbean. Plant Physiology 193: 1197–1212.

53. Liu C, Wang Y, Peng J, Fan B, Xu D, Wu J, Cao Z, Gao Y, Wang X, Li S, et al. 2022. High-quality genome assembly and pan-genome studies facilitate genetic discovery in mung bean and its improvement. Plant Communications 3: 100352.

54. Lowe TM, Eddy SR. 1997. tRNAscan-SE: a program for improved detection of transfer RNA genes in genomic sequence. Nucleic Acids Research 25: 955–964.

55. Ma QH. 2024. Lignin biosynthesis and its diversified roles in disease resistance. Genes 15: 295.

56. Nair R, Schreinemachers P. 2020. Global status and economic importance of mungbean. The mungbean genome 1: 1–8.

57. Nan J, Ling Y, An J, Wang T, Chai M, Fu J, Wang G, Yang C, Yang Y, Han B. 2022. Genome resequencing reveals independent domestication and breeding improvement of naked oat. Gigascience 12: giad061.

58. Nomberg G, Marinov O, Arya GC, Manasherova E, Cohen H. 2022. The key enzymes in the suberin biosynthetic pathway in plants: an update. Plants 11: 392.

59. Ou S, Su W, Liao Y, Chougule K, Agda JRA, Hellinga AJ, Lugo CSB, Elliott TA, Ware D, Peterson T, et al. 2019. Benchmarking transposable element annotation methods for creation of a streamlined, comprehensive pipeline. Genome Biology 20: 275.

60. Pandiyan M, Senthil N, Ramamoorthi N, Muthiah A, Duncan V, Tomooka N, Jayaraj T. 2010. Interspecific hybridization of *Vigna radiata* x 13 wild *Vigna* species for developing MYMV donar. Electronic Journal of Plant Breeding 1: 600–610.

61. Patro R, Duggal G, Love MI, Irizarry RA, Kingsford C. 2017. Salmon provides fast and bias-aware quantification of transcript expression. Nature Methods 14: 417–419.

62. Pertea M, Pertea GM, Antonescu CM, Chang TC, Mendell JT, Salzberg SL. 2015. StringTie enables improved reconstruction of a transcriptome from RNA-seq reads. Nature Biotechnology 33: 290–295.

63. Pollex RL, Hegele RA. 2007. Copy number variation in the human genome and its implications for cardiovascular disease. Circulation 115: 3130–3138.

64. Pryszcz LP, Gabaldon T. 2016. Redundans: an assembly pipeline for highly heterozygous genomes. Nucleic Acids Research 44: e113.

65. Qin S, Fan C, Li X, Li Y, Hu J, Li C, Luo K. 2020. LACCASE14 is required for the deposition of guaiacyl lignin and affects cell wall digestibility in poplar. Biotechnology for Biofuels and Bioproducts 13: 197.

66. Quinlan AR. 2014. BEDTools: the swiss-army tool for genome feature analysis. Curr Protoc Bioinformatics 47: 11.12.11-34.

67. Rhie A, Walenz BP, Koren S, Phillippy AM. 2020. Merqury: reference-free quality, completeness, and phasing assessment for genome assemblies. Genome Biology 21: 245.

68. Robinson JT, Turner D, Durand NC, Thorvaldsdottir H, Mesirov JP, Aiden EL. 2018. Juicebox.js provides a cloud-based visualization system for Hi-C data. Cell Syst 6: 256–258 e251.

69. Scott AJ, Chiang C, Hall IM. 2021. Structural variants are a major source of gene expression differences in humans and often affect multiple nearby genes. Genome Research 31: 2249–2257.

70. Simão FA, Waterhouse RM, Ioannidis P, Kriventseva EV, Zdobnov EM. 2015. BUSCO: assessing genome assembly and annotation completeness with single-copy orthologs. Bioinformatics 31: 3210–3212.

71. Soneson C, Love MI, Robinson MD. 2015. Differential analyses for RNA-seq: transcript-level estimates improve gene-level inferences. F1000Res 4: 1521.

72. Stanke M, Diekhans M, Baertsch R, Haussler D. 2008. Using native and syntenically mapped cDNA alignments to improve de novo gene finding. Bioinformatics 24: 637–644.

73. Sudmant PH, Rausch T, Gardner EJ, Handsaker RE, Abyzov A, Huddleston J, Zhang Y, Ye K, Jun G, Fritz MH, et al. 2015. An integrated map of structural variation in 2,504 human genomes. Nature 526: 75–81.

74. Takahashi Y, Sakai H, Ariga H, Teramoto S, Shimada TL, Eun H, Muto C, Naito K, Tomooka N. 2023. Domesticating *Vigna stipulacea*: chromosome-Level genome assembly reveals VsPSAT1 as a candidate gene decreasing hard-seededness. Frontiers in Plant Science 14: 1119625.

75. Thuan ND. 2011. Expression and inheritance of traits in wild mungbean (Vigna radiata ssp. sublobata) x cultivated mungbean (V. Radiata ssp. radiata) hybrids. James Cook University.

76. Villanueva RAM, Chen ZJ 2019. ggplot2: elegant graphics for data analysis: Taylor & Francis. 160–167.

77. Wang C, Liu C, Roqueiro D, Grimm D, Schwab R, Becker C, Lanz C, Weigel D. 2015. Genome-wide analysis of local chromatin packing in *Arabidopsis thaliana*. Genome Research 25: 246–256.

78. Wang X, Chen L, Ma J. 2019. Genomic introgression through interspecific hybridization counteracts genetic bottleneck during soybean domestication. Genome Biology 20: 1–15.

79. Wang Y, Wang M, Yan X, Chen K, Tian F, Yang X, Cao L, Ruan N, Dang Z, Yin X, et al. 2024. The DEP1 mutation improves stem lodging resistance and biomass saccharification by affecting cell wall biosynthesis in rice. Rice 17: 35.

80. Wu D, Xie L, Sun Y, Huang Y, Jia L, Dong C, Shen E, Ye CY, Qian Q, Fan L. 2023. A syntelog-based pan-genome provides insights into rice domestication and de-domestication. Genome Biology 24: 179.

81. Wu T, Hu E, Xu S, Chen M, Guo P, Dai Z, Feng T, Zhou L, Tang W, Zhan L, et al. 2021. clusterProfiler 4.0: a universal enrichment tool for interpreting omics data. Innovation (Camb) 2: 100141.

82. Xin A, Herburger K. 2021. Precursor biosynthesis regulation of lignin, suberin and cutin. Protoplasma 258: 1171–1178.

83. Xiong W, Wu Z, Liu Y, Li Y, Su K, Bai Z, Guo S, Hu Z, Zhang Z, Bao Y, et al. 2019. Mutation of 4-coumarate: coenzyme a ligase 1 gene affects lignin biosynthesis and increases the cell wall digestibility in maize brown midrib5 mutants. Biotechnology for Biofuels and Bioproducts 12: 82.

84. Xu GC, Xu TJ, Zhu R, Zhang Y, Li SQ, Wang HW, Li JT. 2019. LR_Gapcloser: a tiling path-based gap closer that uses long reads to complete genome assembly. Gigascience 8: giy157.

85. Xu X, Bai G. 2015. Whole-genome resequencing: changing the paradigms of SNP detection, molecular mapping and gene discovery. Molecular Breeding 35: 33.

86. Yang J, Lee SH, Goddard ME, Visscher PM. 2011. GCTA: a tool for genome-wide complex trait analysis. The American Journal of Human Genetics 88: 76–82.

87. Yu G. 2021. Enrichplot: visualization of functional enrichment result. R Package Version 1: 1.

88. Zhang RG, Li GY, Wang XL, Dainat J, Wang ZX, Ou S, Ma Y. 2022. TEsorter: an accurate and fast method to classify LTR-retrotransposons in plant genomes. Horticulture Research 9: uhac017.

89. Zhang T, Peng W, Xiao H, Cao S, Chen Z, Su X, Luo Y, Liu Z, Peng Y, Yang X, et al. 2024. Population genomics highlights structural variations in local adaptation to saline coastal environments in woolly grape. Journal of Integrative Plant Biology 66: 1408–1426.

