## Supplementary Figures for "Telomere-to-telomere, gap-free genome of mung beans (*Vigna radiata*) provides insights into domestication under structural variation"

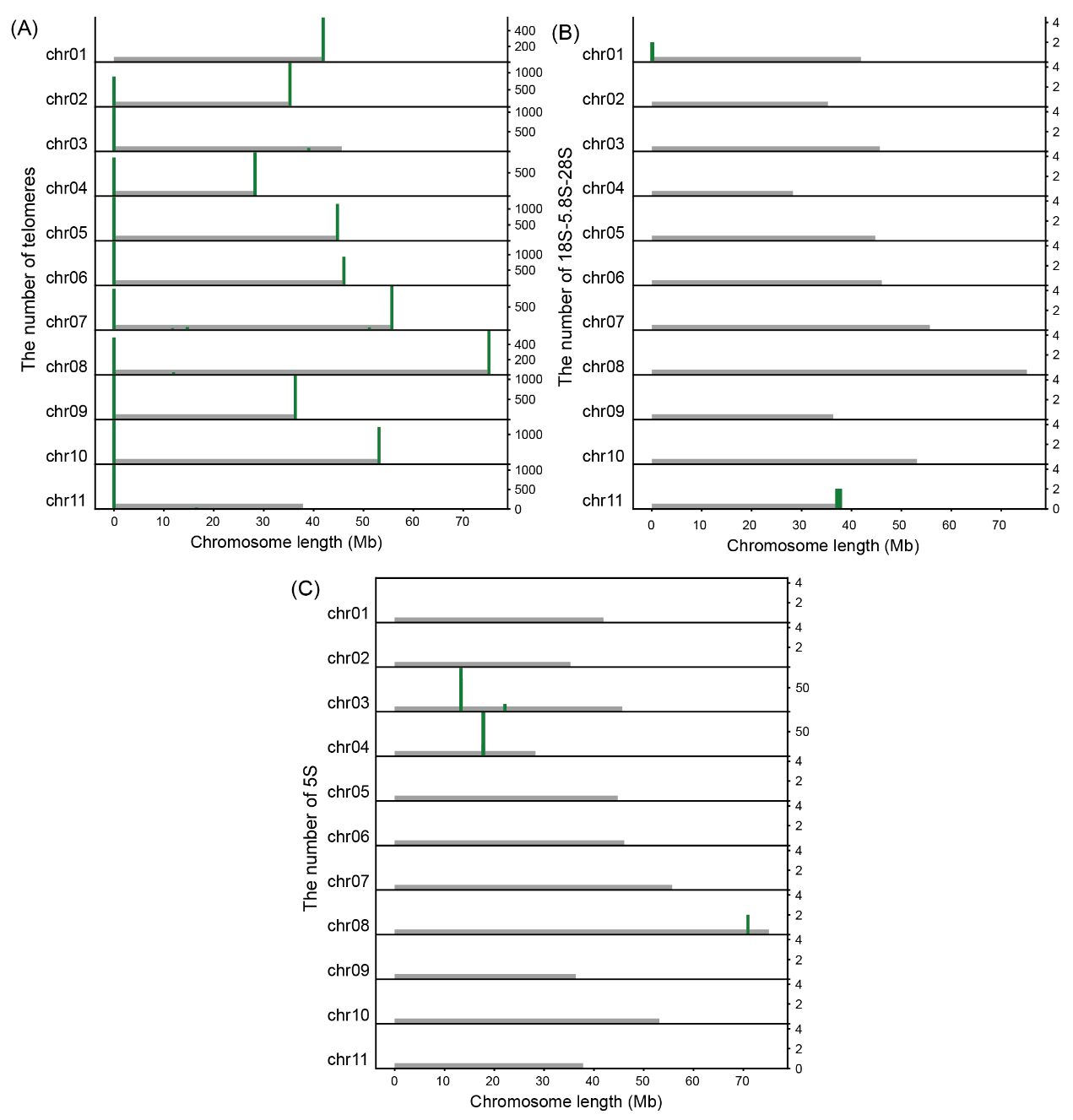
**Supplementary Fig. 1: Characteristics of the Mung Bean Genome.** (A) Distribution and quantity of telomeric sequences across chromosomes. (B) Distribution and quantity of 18S-5.8S-28S rRNA sequences across chromosomes. (C) Distribution and quantity of 5S rRNA sequences across chromosomes.

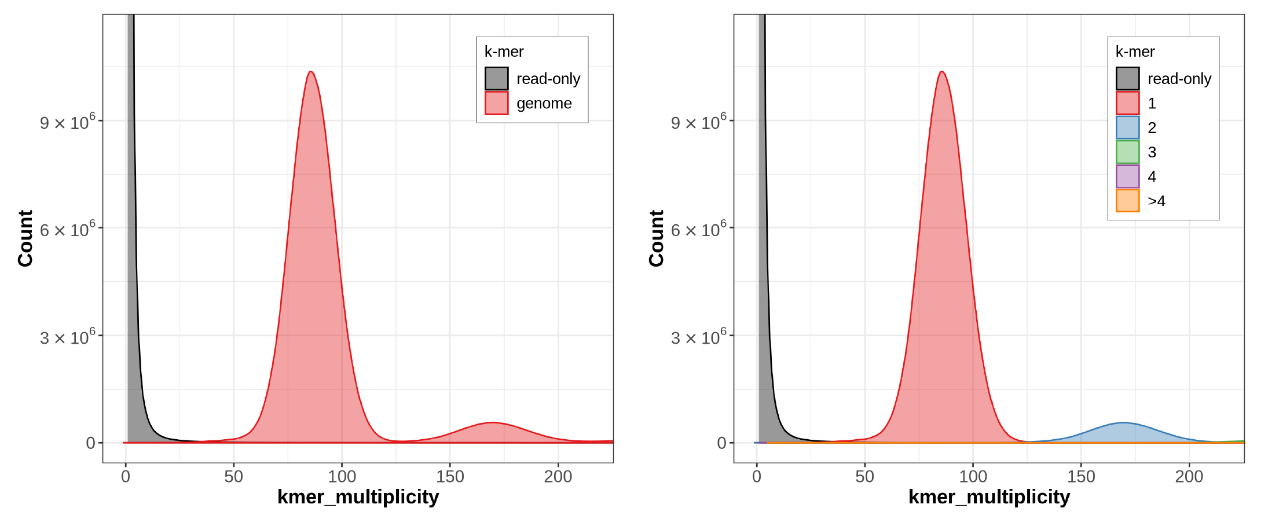
**Supplementary Fig. 2 Genome evaluation using merqury.** Left Panel: K-mer multiplicity distribution comparing read-only k-mers and genome k-mers. The distribution shows the count of k-mers against their multiplicity in the genome. Right Panel: K-mer multiplicity distribution showing the count of k-mers by their multiplicity in read-only k-mers and genome k-mers, color-coded by the number of copies (1, 2, 3, 4, >4).

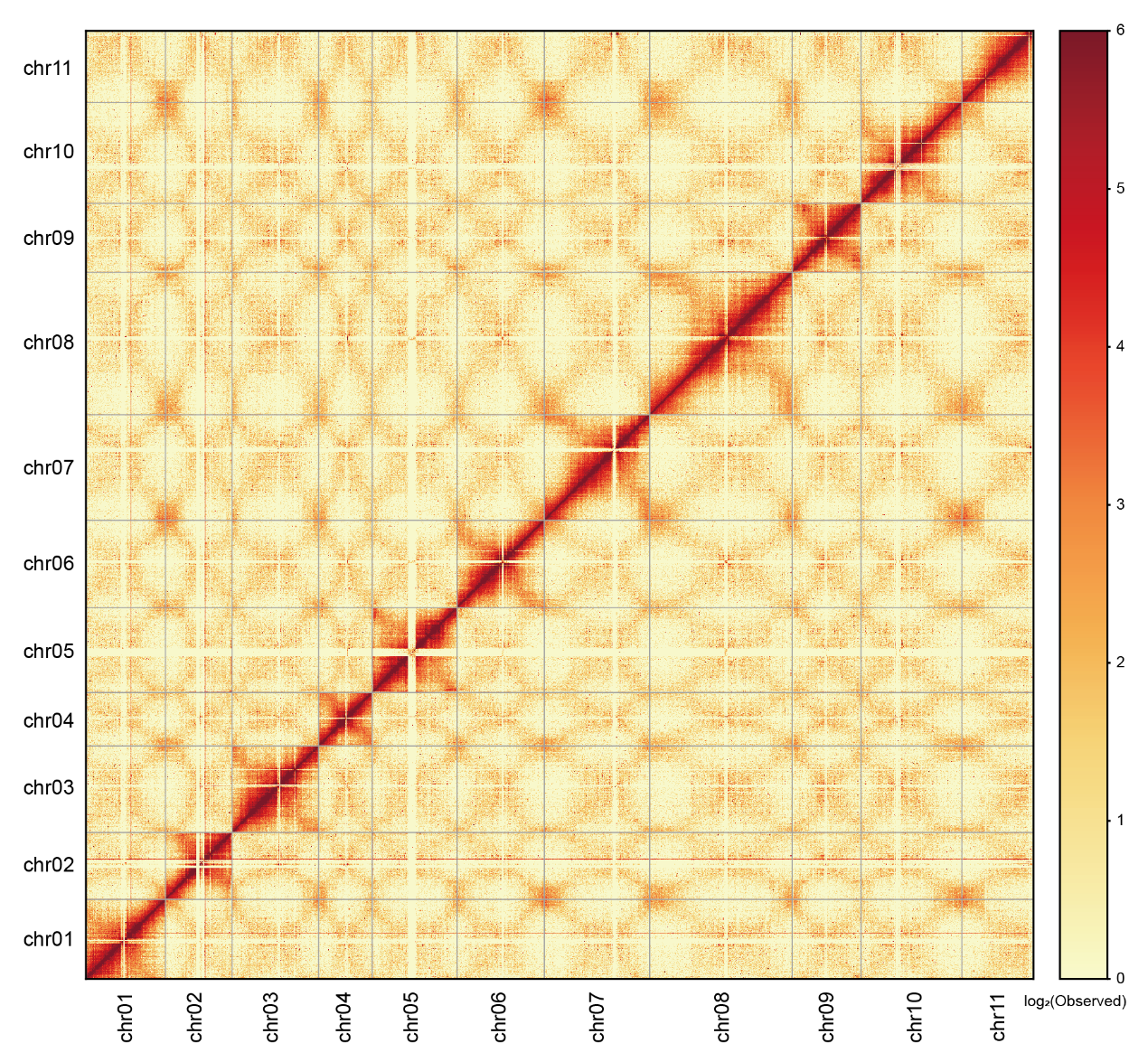

**Supplementary Fig. 3 Hi-C contact map.** This Hi-C contact map displays the chromatin interaction frequencies across the genome. The heatmap shows the intensity of interactions, with the color scale on the right indicating interaction frequency from low (yellow) to high (red). The diagonal line represents interactions within the same regions, while off-diagonal elements indicate interactions between different regions. High-intensity spots along the diagonal suggest regions of high chromatin interaction and potential structural features within the genome.

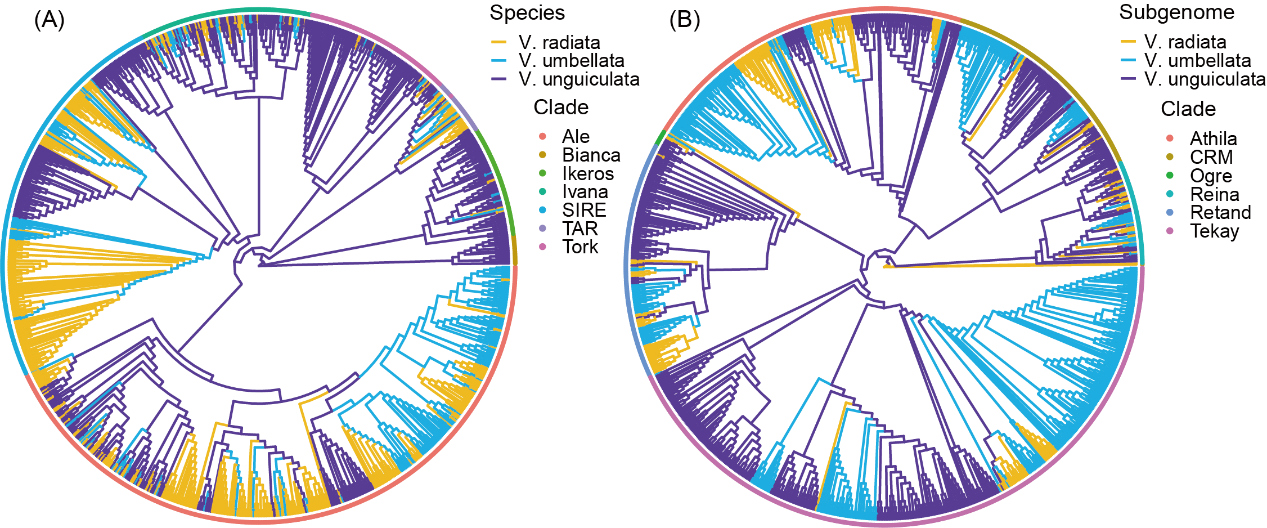

Supplementary Fig. 4 Phylogenetic trees of TE sequences from three genomes.(A) Phylogenetic tree constructed from 1000 randomly selected copia TE sequences from three genomes, showing the evolutionary relationships among the sequences. (B) Phylogenetic tree constructed from 1000 randomly selected gypsy TE sequences from three genomes, showing the evolutionary relationships among the sequences.
